# A closed-loop reinforcement learning framework for rapid compound directed optimization

**DOI:** 10.64898/2026.08.24.745890

**Authors:** Han Wang, Dehua Lu, Weiping Lyu, Siyu Xiu, Cheng Shi, Xiaonan Zhou, Bin Xi, Wei Feng, Yang Xiao, Yanming Chen, Haixian Zhang, Qi Li, Huang Bo, Zhenming Liu

## Abstract

Generative artificial intelligence (AI) holds transformative potential for drug discovery, yet existing architectures typically operate in open loops without experimental feedback. Here we introduce rapid compound directed optimization (RCDO), a closed-loop reinforcement learning framework that accelerates the optimization process by bridging dry-lab computation with wet-lab feedback. RCDO couples a three-dimensional structure-guided generative model with a multi-level reward system updated after each design cycle using experimental measurements from all synthesized compounds, including inactive or developability-failed compounds. By continuously aligning the generative model with accumulated wet-lab measurements, RCDO substantially compresses optimization timelines. We evaluated RCDO through retrospective benchmarking against historical optimization trajectories and prospective wet-lab campaigns targeting ROR1, NLRP3, and NSD3. Across prospective evaluations, RCDO rapidly resolved key optimization bottlenecks within two to three design cycles: improving the oral exposure of an ROR1 inhibitor by 40-fold while maintaining antitumor efficacy, reducing CYP2C19 inhibition of an NLRP3 antagonist by 20-fold while preserving inflammasome activity, and boosting the binding affinity of an NSD3 hit by 18-fold. By directly coupling wet-lab feedback to generative learning, RCDO establishes an efficient platform for compound directed optimization, transforming AI-driven drug discovery from static generation into continuous experimental adaptation.

## 1. Introduction

Transforming initial bioactive hits into developable drug candidates remains one of the most challenging stages in drug discovery. A compound with strong target activity is rarely sufficient for therapeutic development, as successful candidates must simultaneously achieve balanced potency, selectivity, pharmacokinetic properties, and safety profiles. Achieving this balance requires iterative design-make-test-analyze (DMTA) cycles, in which medicinal chemists continuously learn from experimental measurements and progressively refine molecular structures. Thus, molecular optimization is fundamentally an adaptive evolutionary process driven by accumulated experimental knowledge.

Recent advances in artificial intelligence (AI) have expanded the scope of computational molecular design. Generative models can explore chemical space, incorporate predefined design objectives and propose compounds for experimental evaluation. Yet many AI-driven discovery workflows remain open-loop without experimental feedback: molecules are generated computationally and then tested experimentally, with limited feedback from these measurements to the generative model. Unfavorable results and unsuccessful designs can reveal important constraints, but this information is often not used to update subsequent molecular generation. This separation limits the capacity of generative models to learn continuously during a medicinal chemistry campaign. To clarify the distinctions among molecular design paradigms, Figure 1A schematically compares how optimization and experimental feedback are incorporated into their workflows.

**Figure 1.**
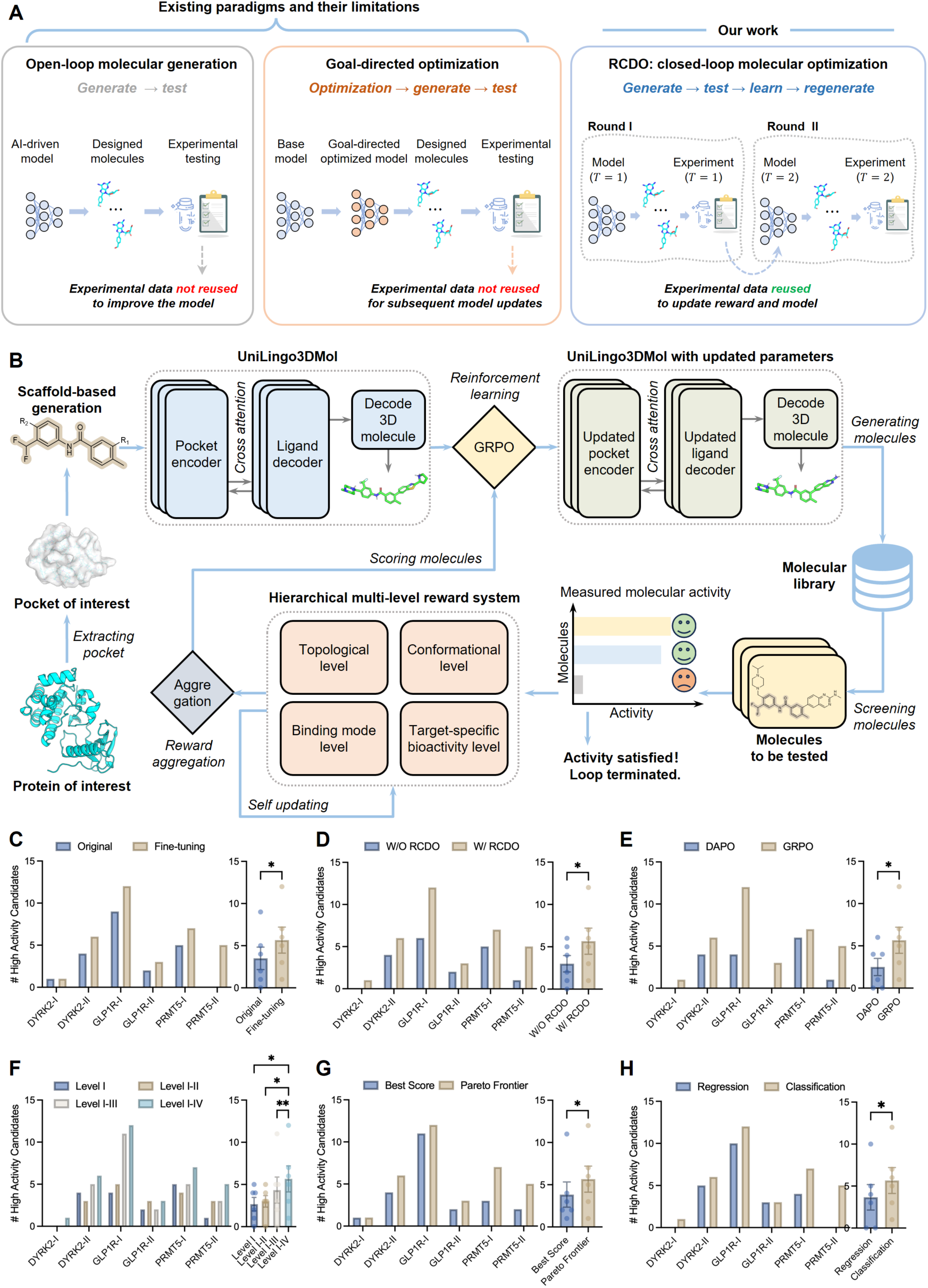
Architecture of the RCDO framework and ablation studies used to select its optimal configuration. **(A)** Schematic comparison of molecular design paradigms. In open-loop molecular generation, candidate compounds are generated and experimentally tested, but resulting measurements are not used to update the generative model. In goal-directed optimization, predefined computational objectives guide molecular generation, while experimental measurements are not incorporated into subsequent model updates. In RCDO, experimental measurements from each cycle are used to update reward and model, enabling closed-loop adaptation of subsequent molecular designs. **(B)** Schematic of the RCDO framework. A protein of interest is first processed to extract its binding pocket, which is then used to condition scaffold-based 3D molecular generation via UniLingo3DMol. RCDO employs a hierarchical multi-level reward system that includes topological, conformational, binding mode, and target specific bioactivity levels to evaluate and score the generated molecules. These scores are subsequently aggregated to update the original UniLingo3DMol model through GRPO based reinforcement learning. Iterative cycles of generation, evaluation, and model updating progressively expand the molecular library. The process continues until the desired activity criteria are achieved. **(C–H)** Six ablation experiments, each isolating a single design choice across the six scaffolds and three targets. In each panel, the y-axis represents the number of high-activity compounds (defined as those with an ECFP Tanimoto similarity greater than 0.9 to their corresponding historical hit). The left bars show counts per scaffold, while the right bars present the pooled results across scaffolds (mean ± SEM, with individual scaffolds overlaid as points). Statistical significance of the pooled comparison is indicated above the summary bars (*P < 0.05; **P < 0.01; ns, not significant). **(C)** The Fine-tuned versus original UniLingo3DMol. **(D)** The complete RCDO directed optimization loop versus a baseline without RCDO. **(E)** The GRPO versus DAPO model updating algorithm. **(F)** The cumulative reward hierarchy at increasing depth. **(G)** The Pareto-frontier versus best-score recommendation. **(H)** The classification-based versus regression-based QSAR model for the bioactivity reward.

Existing computational methods address complementary aspects of molecular optimization. Structure-based three-dimensional (3D) generative models, including TargetDiff^1^, Lingo3DMol^2^, MolCRAFT^3^, and related approaches^4^, generate molecules conditioned on protein binding pockets while explicitly representing molecular geometry. However, these models are designed primarily to generate candidate structures rather than to learn iteratively from successive rounds of experimental measurements. Ligand-based reinforcement learning (RL) methods, such as REINVENT4^5^, enable goal-directed optimization but generally operate on molecular representations that do not explicitly encode 3D protein-ligand interactions. Recent work has incorporated RL into 3D molecular generation. SeFMol^6^ formulates pocket-conditioned diffusion denoising as a Markov decision process, enabling RL-guided adjustment of molecular structures during 3D generation. Although these methods can adapt individual molecules during computational generation, their generative models remain fixed across experimental cycles rather than evolving in response to new measurements.

This limitation is particularly restrictive in small molecule optimization, where each DMTA cycle can reveal new constraints and redirect subsequent design. Such constraints are often difficult to predict before compounds are synthesized and tested. A key unmet challenge in AI-driven drug discovery is therefore to develop a framework that bridges dry-lab computation with wet-lab feedback to continuously adapt generative models across successive DMTA cycles.

Here, we introduce rapid compound directed optimization (RCDO), a closed-loop reinforcement learning framework that accelerates the optimization process by bridging dry-lab computation with wet-lab feedback. RCDO couples a 3D structure-guided generative model, UniLingo3DMol^7^, with a multi-level reward system organized into four hierarchical levels: topological validity, conformational plausibility, binding mode complementarity, and target-specific bioactivity. After each DMTA cycle, experimental measurements from all synthesized compounds, including inactive or developability-failed compounds, update the reward functions at the corresponding levels, and the generative model is then refined by RL against updated reward functions. Both the reward functions and the generator thus evolve with accumulated wet-lab measurements, allowing subsequent DMTA cycles to sample from a distribution already shaped by experimental measurements rather than repeatedly proposing compounds that downstream filters would reject.

We optimized the RCDO framework through retrospective benchmarking against historical optimization trajectories, then evaluated it in prospective wet-lab optimization campaigns targeting the kinase-like receptor ROR1, the NLRP3 inflammasome, and the NSD3 PWWP1 domain. Across these prospective evaluations, RCDO rapidly resolved key optimization bottlenecks within two to three DMTA cycles: improving the oral exposure of an ROR1 inhibitor by 40-fold while maintaining antitumor efficacy, reducing CYP2C19 inhibition of an NLRP3 antagonist by 20-fold while preserving inflammasome activity, and boosting the binding affinity of an NSD3 hit by 18-fold. Together, these results show that RCDO can support rapid compound directed optimization across distinct discovery challenges and provide an experimental feedback-driven framework for AI-assisted molecular evolution.

## 2. Results

### 2.1. Benchmarking of The RCDO Framework

Before any prospective deployment, we optimized the RCDO framework (Figure 1B) *in silico* to select its architecture and parameters. RCDO uses UniLingo3DMol^7^ as its base generator to propose 3D molecular candidates from an input seed molecule and target-binding pocket. The generated candidates are evaluated using a hierarchical reward system, after which RL updates the generative model and Pareto-frontier selection identifies candidates for the next optimization cycle. We benchmarked the framework using three historical optimization trajectories curated from previous studies^8–10^.

The benchmark set comprised three historical optimization trajectories spanning distinct target classes: the kinase DYRK2^8^, the membrane receptor GLP1R^9^ and the methyltransferase PRMT5^0^. Each target contributed two completed optimization scaffolds, suffixed I and II, yielding six benchmark tasks in total (Figure S1A-C). Each task contained a historical starting hit and a set of experimentally validated active analogs. We initialized RCDO with the starting hit and assessed whether it could recover structures resembling the validated analogs without access to their downstream experimental data. This retrospective design enabled direct comparison between generated candidates and experimentally validated optimization outcomes. For each task, we scored each configuration by the number of high-activity candidates among the top 200 molecules ranked by the recommendation strategy. For this retrospective benchmark, we operationally defined a high-activity candidate as a generated molecule with an ECFP Tanimoto similarity greater than 0.9 to at least one experimentally validated active analog. This definition provided a structural-rediscovery proxy for bioactivity rather than direct evidence of biological activity. Although high structural similarity to a validated active analog may indicate likely activity under the molecular similarity principle, this proxy may fail near activity cliffs or overlook active molecules with distinct structures.

We optimized the RCDO architecture through six ablations, each isolating one key design choice on the benchmark (Figure 1C-H). Together, the six choices trace the path a seed molecule follows from generation to recommendation: (i) the molecular tokenization and fine-tuning of the UniLingo3DMol base generator, which sets the chemical edits the model can propose; (ii) the contribution of the complete RCDO directed optimization loop, which couples the generator to reinforcement learning; (iii) the policy-update algorithm, which converts reward into model updates; (iv) the depth of the multi-level reward hierarchy, which defines what the framework treats as an improved molecule; (v) the molecular recommendation strategy, which decides which candidates advance; and (vi) the formulation of the bioactivity reward, which turns predicted activity into a learning signal.

The first two ablations evaluated the foundational choices. The fine-tuned UniLingo3DMol model with revised tokenization obtained more high-activity candidates than the original model (Figure 1C). The revised tokenization improved the model’s ability to generate more localized modifications, which were poorly represented by the original tokenization. The complete RCDO directed-optimization loop also nominated more high-activity candidates than a baseline without RCDO (Figure 1D). The remaining four ablations evaluated individual components of the optimization loop. For model updating strategy, group relative policy optimization (GRPO) outperformed the competing DAPO algorithm (Figure 1E). For reward design, the complete four-level reward hierarchy (levels I-IV) outperformed variants truncated at Level I, Level I-II, and Level I-III (Figure 1F). The complete hierarchy integrated topological validity, conformational plausibility, binding mode complementarity, and target-specific bioactivity. For candidate recommendation, the Pareto-frontier strategy outperformed best-score selection (Figure 1G). By retaining molecules that were optimal along at least one reward dimension, this strategy preserved a chemically diverse set of complementary candidates. For the bioactivity reward, the classification-based quantitative structure–activity relationship (QSAR) model identified more high-activity candidates than the regression-based model (Figure 1H, and Figure S1D-F). The benchmark activity data were derived from heterogeneous assay formats and showed skewed value distributions. Under these conditions, the regression-based model generated noisy, poorly calibrated reward signals that destabilized model updates. By contrast, the classification-based model provided a more robust reward signal and enriched for experimentally active chemotypes. These results indicate that matching the reward formulation to the quality and distribution of the underlying data, rather than defaulting to regression, improved learning across heterogeneous benchmarks. Based on these ablations, the configuration selected for prospective evaluation comprised the fine-tuned base model, the complete RCDO loop, GRPO, four-level reward hierarchy, Pareto-frontier recommendation, and classification-based QSAR.

Moreover, we also compare RCDO with REINVENT4^5^, a state-of-the-art 2D reinforcement learning framework for generative molecule design, on the same six benchmark scaffolds (Figure S1A-C). REINVENT4 was configured with the LibInvent generator and the officially released pretrained weights, optimized via the default policy using a scoring function that combined the official 2D constraint components with an AutoDock-GPU^11^ docking reward, and the top 200 molecular candidates were generated by ranking the molecules by their docking scores. Under our structural-rediscovery metric, REINVENT4 nominated zero high-activity candidates across all six tasks, whereas RCDO recovered validated analogs on every scaffold (Figure S1G). RCDO avoids this failure mode through two design principles absent in REINVENT4. First, the Level I–IV hierarchical reward injects 3D binding-mode complementarity (Level III) as an intermediate learning signal, providing denser gradient information than a single scalar docking score and guiding the generator toward the correct substituent edits required to maintain or improve the binding pose. Second, the Pareto-frontier recommendation strategy avoids the unreasonable molecular structures that result from relying solely on docking scores.

Under the configured framework, RCDO produced high-activity candidates across the three historical trajectories. This collective recovery, read directly from the number of high-activity candidates, validated both the fine-tuned generator and the multi-level reward system on independent retrospective benchmarks. Having fixed these parameters, we next deployed the same configuration prospectively to three live medicinal chemistry campaigns.

### 2.2. RCDO-Guided Optimization of Oral Exposure of ROR1 Inhibitors for Triple-Negative Breast Cancer

#### 2.2.1. Pharmacokinetic Liabilities and Metabolic Profiling of Hit Compound 59

Triple-negative breast cancer (TNBC) lacks broadly effective targeted treatment options, and ROR1 has emerged as a therapeutically relevant kinase-like receptor in this disease context^12,13^. In TNBC cells, ROR1-mediated signaling is linked to PI3K/AKT and STAT3/NF-κB pathway activation, which supports proliferation, survival, migration, and tumor development^14–16^ (Figure 2A). A previous structure-based campaign from marketed drug ponatinib yielded Compound **59** (Figure 2B, and Figure S2D) a potent ROR1 inhibitor with anti-TNBC activity (K_D_ = 0.058 μM, IC_50_ MDA-MB-231 = 0.086 μM), but its further development was limited by poor oral exposure^17^ (F = 3.51%, AUC_last 0-8h_ = 36.4 h*ng/mL) (Table S1). Developability profiling identified two liabilities that could explain this behavior: low membrane permeability (LogPe = -6.59) (Figure S2A) and rapid human liver microsomal turnover (T_1/2_ = 12.8 minutes) (Figure S2B). The supplementary analysis further linked these liabilities to highly polar groups, including the piperazine ring and 2-aminoquinazoline, and SMARTCyp nominated methylene and methyl groups around the piperazine nitrogen atoms, together with the methyl group para to the benzamide moiety, as likely metabolic hotspots (Figure S2C). We therefore used RCDO to reduce polarity and modify labile metabolic sites while maintaining ROR1 binding and cellular anti-TNBC activity.

**Figure 2.**
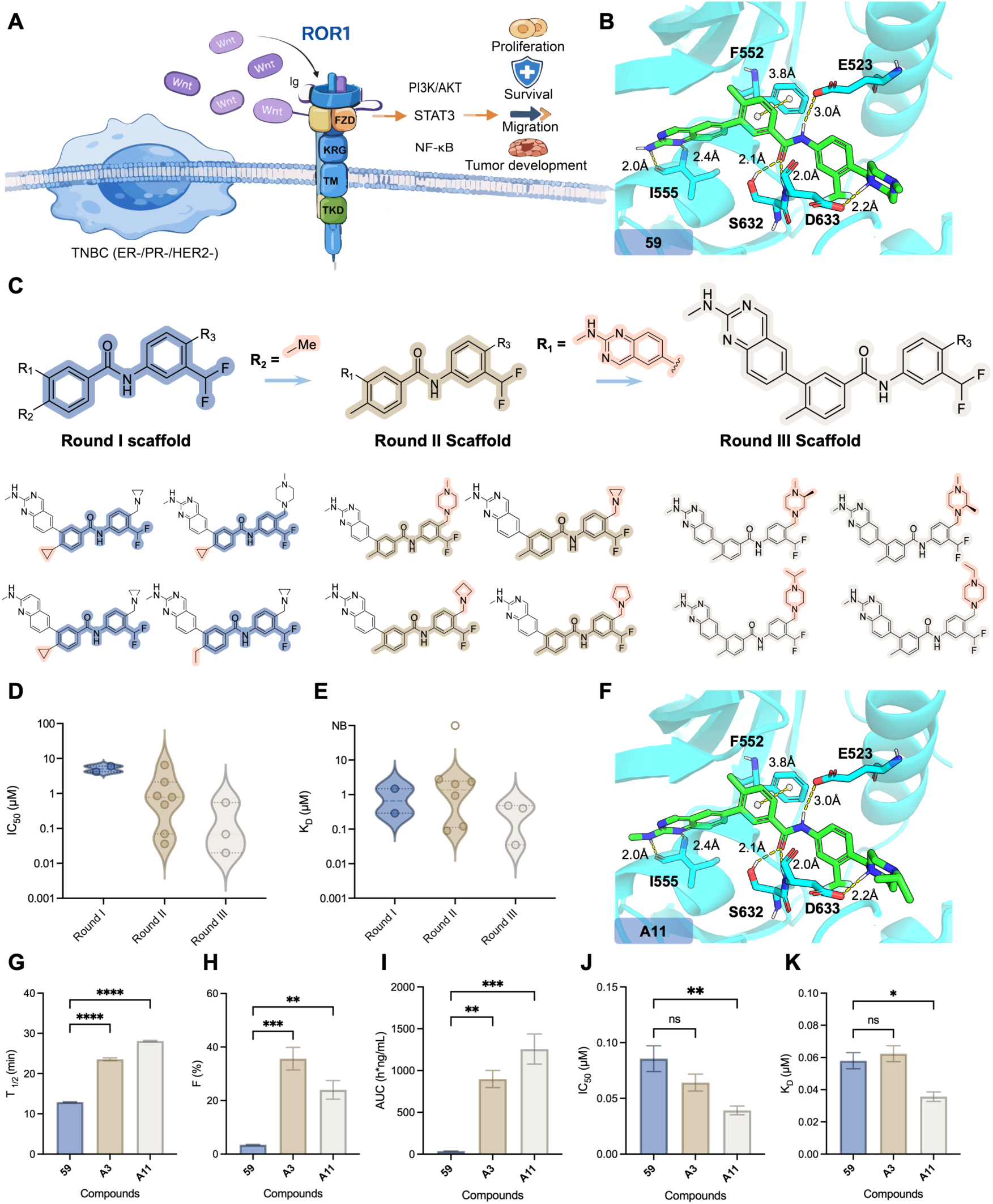
RCDO-guided optimization of ROR1 inhibitors for improved oral exposure and retained potency. **(A)** Schematic illustration of the ROR1-mediated signaling pathway in TNBC cells. **(B)** Binding mode of Compound **59** in the ROR1 binding pocket, showing key interacting residues and intermolecular distances. **(C)** Visualization of scaffold evolution from Round I to Round III under the RCDO framework, highlighting the progressive structural changes across the three optimization cycles. **(D-E)** Distributions of IC_50_ values in MDA-MB-231 cells and ROR1 binding affinity K_D_ values for compounds from Round I, II, and III scaffolds. **(F)** Binding mode of the optimized Compound **A11**. **(G-K)** Stepwise comparison of the starting Compound **59**, intermediate Compound **A3** and optimized Compound **A11**. **(G)** Liver microsome metabolic stability, expressed as half-life T_1/2_ (T_1/2_, minute, n = 3). **(H)** Oral bioavailability (F, %, n = 3). **(I)** Area under the plasma concentration-time curve (AUC), representing total systemic exposure to the compound following oral administration (AUC, h*ng/mL, n = 3). **(J)** IC_50_ MDA-MB-231 values (μM, n = 3). **(K)** ROR1 binding affinity K_D_ values. Chemical structures of the three compounds are provided in Figure S2D. Data are presented as mean ± SEM or individual data points. Statistical significance was assessed using ordinary one-way ANOVA: ns (not significant), *p < 0.05, **p < 0.01, ***p < 0.001, ****p < 0.0001.

#### 2.2.2. Iterative Structural Optimization and Discovery of Compound A11

We next applied the RCDO framework to the scaffold of Compound **59** via three iterative DMTA cycles, following the scaffold-evolution workflow illustrated in Figure 2C. In the first generation round, the ROR1 binding pocket and the three-dimensional conformation of Compound **59** served as seed templates for molecular generation. To constrain unreasonable lipophilicity and rationalize oral druggability optimization, quantitative structure-activity relationship (QSAR) models were integrated into this multi-layer reward system (Table S2, and Figure S2E-H).

For the first round, cyclopropyl and ethyl substituents frequently appeared at the R_2_ position among the top four highest-ranked generated compounds (Figure 2C, left). To characterize the functional impact of R_2_ modification, we fixed all other core fragments and synthesized two analogs **A1** and **A2** with cyclopropyl and ethyl groups installed at the R_2_ site, respectively (Table S3). Biological validation revealed that replacing the native methyl group at R_2_ with cyclopropyl or ethyl reduced ROR1 binding affinity. Accordingly, the R_2_ substituent was locked as methyl for all molecular generation tasks in the second round. Prior to launching the second generation cycle, we imported all activity data collected from first-round compounds into the reward system to refine the model’s predictive performance. The top four prioritized molecules from Round II were dominated by structural variations at the R_3_ position (Figure 2C, middle); therefore, we fixed all other scaffold moieties and synthesized analogs **A4**–**A10** bearing diversified R_3_ substituents for experimental validation (Table S4). Notably, most high-scoring compounds generated in both Round I and Round II featured substitution of the original 2-aminoquinazoline R_1_ group with 2-(methylamino)quinazoline. To explicitly evaluate the biological performance of this R_1_ modification, we specially synthesized analog **A3** carrying the 2-(methylamino)quinazoline moiety at R_1_. Biological profiling demonstrated that **A3** exhibited robust ROR1 binding affinity and potent antiproliferative activity against TNBC cells (K_D_ = 0.062 μM, IC_50_ MDA-MB-231 = 0.064 μM) (Figure S2D). We further quantified its oral bioavailability in mice: **A3** yielded an oral bioavailability value (F = 35.64%), representing a nearly 10-fold elevation relative to the parent Compound **59** (F = 3.51%) (Table S1). Given that oral systemic exposure (AUC) is a pivotal pharmacokinetic endpoint for judging oral drug feasibility, further optimization of **A3**’s oral bioavailability was still required.

To strengthen the model’s capacity to optimize oral druggability, we experimentally measured the membrane permeability and liver microsomal metabolic stability of all second-round compounds (Table S4). The resulting experimental datasets were utilized to train dedicated QSAR models, which were subsequently incorporated into the reward system to drive third-cycle molecular generation (Figure S2I-J). Considering that the introduction of 2-(methylamino)quinazoline at R_1_ in **A3** substantially improved oral bioavailability, we fixed the R_1_ substituent as 2-(methylamino)quinazoline throughout the third generation workflow, followed by chemical synthesis of analogs **A11**–**A13** (Table S5). Among these candidates, analog **A11** displayed prominent ROR1 binding affinity and strong anti-TNBC proliferative activity (K_D_ = 0.035 μM, IC_50_ MDA-MB-231 = 0.039 μM) (Figure S2D). Subsequent pharmacokinetic assays confirmed that **A11** achieved markedly enhanced oral systemic exposure (AUC) compared with **A3**, making it a promising candidate for oral antitumor therapy (Table S1).

Collectively, we exploited the RCDO framework to iteratively optimize the oral bioavailability of starting Compound **59** across three successive DMTA cycles. Across all three rounds of experiments, the synthetic analogs exhibited an overall trend of gradual elevation in both their binding affinity for ROR1 and anti-proliferative activity against TNBC (Figure 2D-E). The representative hits from Round II (**A3**) and Round III (**A11**) achieved remarkable improvements over parental Compound **59** across all core pharmacokinetic and biological readouts, including liver microsomal metabolic stability, oral bioavailability, systemic oral exposure, ROR1 binding affinity, and anti-TNBC proliferative activity (Figure 2G-K). Collectively, these data reveal progressive property optimization of ROR1 inhibitors through iterative modification cycles, confirming the capability of RCDO in directed molecular evolution.

The final lead compound **A11** not only retains potent in vitro ROR1 binding and anti-TNBC proliferative activity, but also possesses drastically improved druggability. Moreover, **A11** preserved the canonical ROR1 pocket interaction pattern (Figure 2F), while presenting a superior developability profile compared with parent Compound **59** (Figure 2B).

#### 2.2.3. *In Vitro* and *In Vivo* Anti-TNBC Activity Evaluation of Compound A11

We then evaluated whether Compound **A11** retained cellular target engagement and anti-TNBC activity in the assays summarized in Figure 3. In kinase profiling across 360 assays, **A11** showed only 12 interactions at 0.5 μM, demonstrating a favorable kinase selectivity profile under the tested conditions (Figure 3A). In MDA-MB-231 cells, **A11** reduced ROR1 phosphorylation and shifted apoptosis-associated markers toward a pro-apoptotic state, with increased Bax, reduced Bcl-2, and induction of cleaved PARP (Figure 3B). **A11** also suppressed colony formation across the tested concentration range and slowed scratch-wound closure over 12-48 hours, consistent with reduced proliferative and migratory capacity in vitro (Figure 3C-D). In a subcutaneous MDA-MB-231 xenograft model, oral **A11** reduced tumor growth more effectively than the parent Compound **59**, with endpoint tumor-volume differences evident on day 27 (Figure 3E). In an orthotopic MDA-MB-231-luciferase model, oral dosing at 20 mg/kg reduced tumor-associated bioluminescence between days 7 and 43 and lowered ex vivo fluorescence signals in major organs at the end of treatment (Figure 3F). Given the prominent oral anti-TNBC activity of compound **A11**, we performed an acute toxicity assay to evaluate its safety profile. The results showed that no mortality or body weight loss was observed in mice throughout the 14-day observation period after a single dose of 1000 mg/kg (Figure S3A-C).

**Figure 3.**
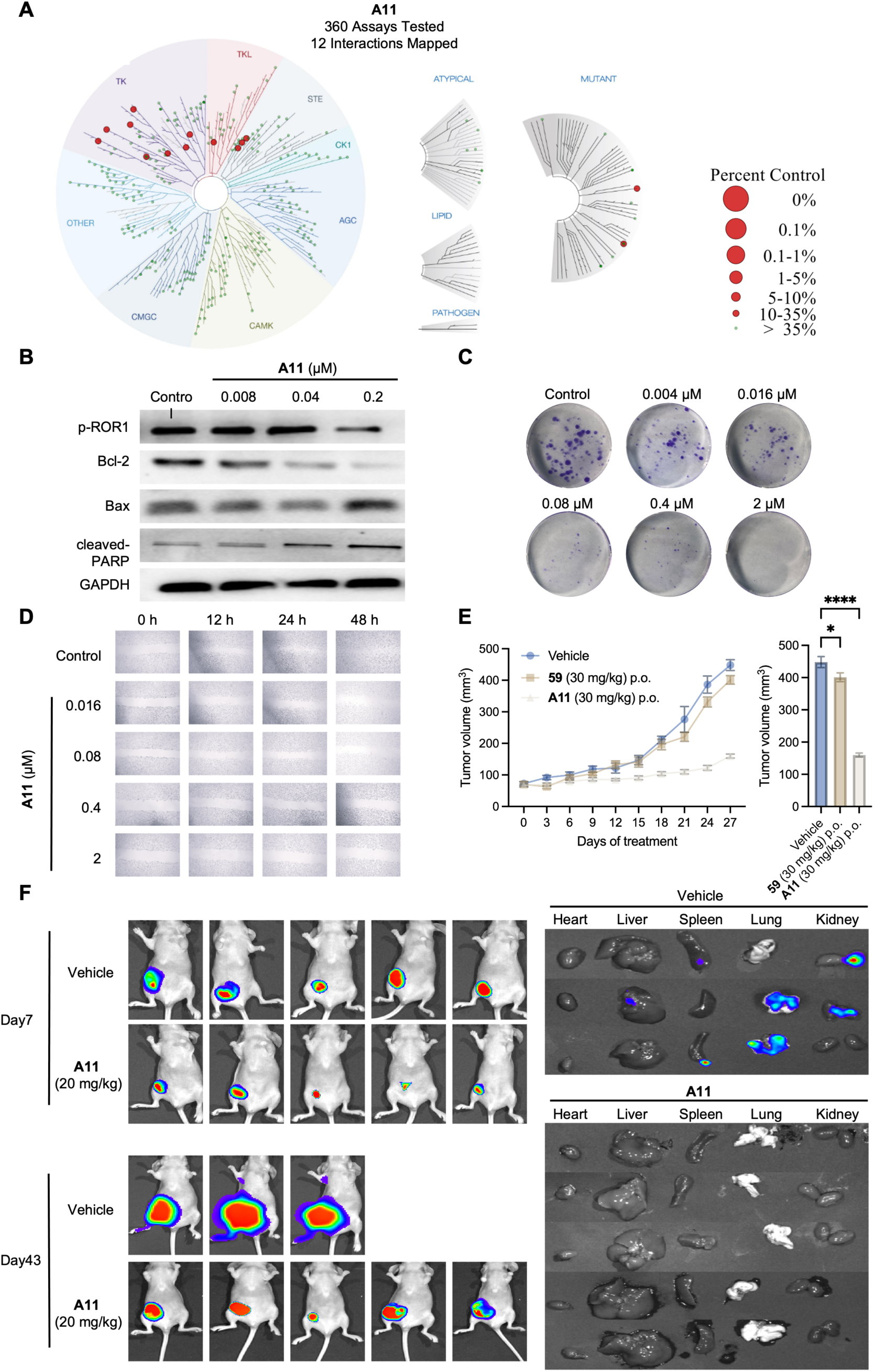
*In vitro* and *in vivo* antitumor activity of Compound A11. **(A)** Kinase selectivity profile of Compound **A11** tested at 0.5 μM against a panel of 360 kinase assays. **(B)** Inhibition of ROR1-mediated signaling pathways by Compound **A11**. **(C-D)** Inhibition of proliferation and migration in MDA-MB-231 cells by Compound **A11**. **(C)** Effect of Compound **A11** on colony formation of MDA-MB-231 cells. **(D)** Effect of Compound **A11** on scratch wound healing of MDA-MB-231 cells. **(E)** Tumor growth curves and endpoint tumor volumes under different treatments. The left panel shows tumor volume (mm^3^) measured over time for the Vehicle, Compounds **59**, and **A11** groups, while the right panel shows tumor volume for each group on day 27. Data are presented as mean ± SEM. Statistical significance was assessed using ordinary one-way ANOVA: ns (not significant), *p < 0.05, **p < 0.01, ***p < 0.001, ****p < 0.0001. **(F)** Compound **A11** inhibited tumor proliferation and metastasis in an orthotopic breast cancer model. An orthotopic xenograft model was established using luciferase-expressing MDA-MB-231 cells, and treatment was administered every three days. *In vivo* imaging was performed on days 7 and 43 after treatment initiation. After 43 days of treatment, the mice were sacrificed, and major organs were collected for fluorescence imaging.

Together, the ROR1 campaign shows how RCDO can couple structure-guided generation with round-by-round experimental feedback to improve a parent inhibitor across potency, permeability, metabolic stability, and oral anti-TNBC activity. These findings demonstrate that the RCDO framework possesses robust directed optimization capabilities, enabling the improvement of pharmaceutical properties of drug molecules while preserving their target bioactivity.

### 2.3. RCDO-Guided Optimization of Metabolic Stability of SN3-1 for NLRP3 Inflammasome Inhibition

#### 2.3.1. Pharmacokinetic Liabilities and Metabolic Profiling of SN3-1

The NLRP3 inflammasome is a central innate immune signaling platform that converts cellular danger sensing into caspase-1 activation, IL-1β maturation and inflammatory tissue injury. Dysregulated NLRP3 activation contributes to sterile inflammatory diseases, gout and inflammatory bowel disease, making this pathway an attractive target for small-molecule intervention (Figure 4A). A useful NLRP3 inhibitor, however, must do more than block IL-1β release in cells. It must retain target engagement while avoiding metabolic liabilities, drug-drug interaction risk and tissue toxicity during repeated dosing. This balance has been difficult to achieve. **MCC950** established pharmacological proof of concept for selective NLRP3 inhibition, but the broader sulfonylurea chemotype has been associated with development-limiting safety concerns. Compound **SN3-1** (Figure 4B), derived from the molecular scaffold of **MCC950** in our group’s earlier work^18^, was designed as a potent NLRP3 inhibitor and exhibited promising anti-inflammatory activity. Nevertheless, its translational potential remains restricted by unaddressed developability risks. **SN3-1** was selected as the starting point for NLRP3 optimization because it combined strong cellular potency with direct target binding. In LPS-primed macrophage assays activated with nigericin, **SN3-1** inhibited IL-1β release with an IC_50_ of 8 nM, and prior binding evaluation showed direct NLRP3 engagement with a K_D_ of 6.9 nM (Figure 4C). These data established **SN3-1** as a credible parent inhibitor for lead optimization.

**Figure 4.**
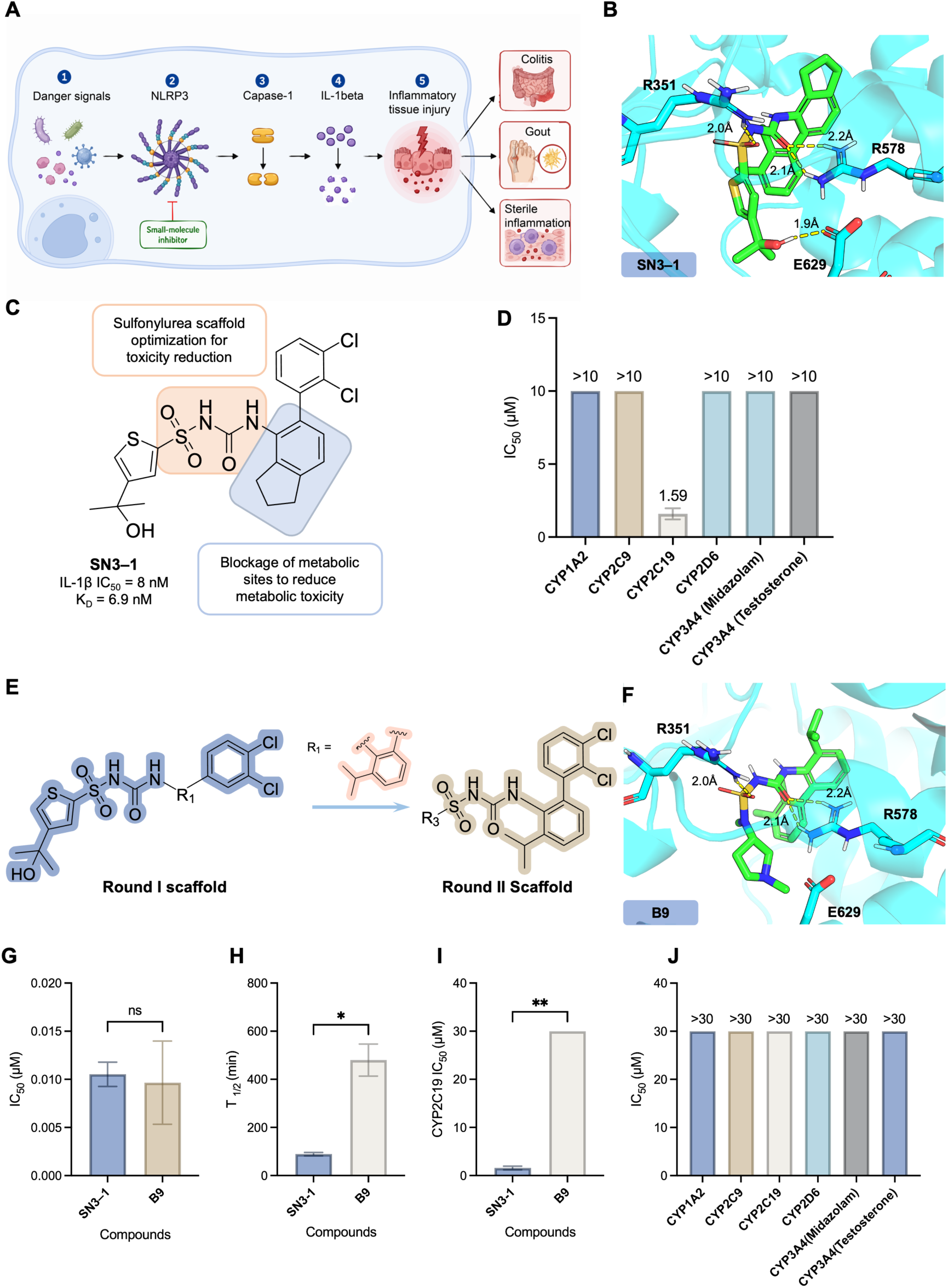
RCDO-guided optimization of NLRP3 inhibitors for improved metabolic stability and reduced CYP inhibition. **(A)** Schematic illustration of the NLRP3 inflammasome activation pathway. **(B)** Binding mode of Compound **SN3-1** within the NLRP3 binding pocket, showing key interactions with surrounding residues. **(C)** Chemical structure of Compound **SN3-1** and the structure-guided optimization strategy of its sulfonylurea scaffold. Optimization aimed to reduce potential toxicity and block metabolically labile sites while preserving potent NLRP3 inhibition. **(D)** Inhibitory activity of **SN3-1** against major human cytochrome P450 (CYP) isoforms, revealing preferential inhibition of CYP2C19. **(E)** Visualization of RCDO-guided scaffold evolution from Round I to Round II scaffold, highlighting structural remodeling intended to improve metabolic and safety-related properties. **(F)** Binding mode of the optimized Compound **B9** in the NLRP3 binding pocket. **(G-J)** Comparison of Compounds **SN3-1** and **B9** during stepwise optimization. **(G)** Cellular NLRP3 inhibitory activity, expressed as IC_50_. **(H)** Liver microsome metabolic stability, expressed as half-life T_1/2_. **(I)** CYP2C19 inhibitory activity comparison among the representative compounds. **(J)** CYP inhibition profile of the optimized Compound **B9** across major human CYP isoforms. Data are presented as mean ± SEM or individual data points where applicable. Statistical significance was assessed as indicated in the figure: ns, not significant; ***p < 0.001. Statistical significance was assessed using two-sided Student’s t-tests: ns (not significant), *p < 0.05, **p < 0.01, ***p < 0.001, ****p < 0.0001.

Developability profiling revealed that Compound **SN3-1** still carried important liabilities. In a 12-day repeat-dose mouse study, **SN3-1** produced liver pathology across tested dose groups. Compared with the control liver, the 30 mg/kg group showed focal hepatocellular necrosis with inflammatory infiltration, whereas 100 and 300 mg/kg dosing produced more evident hepatocellular vacuolar degeneration (Figure S4A). These findings indicated that high NLRP3 potency did not eliminate scaffold-associated liver risk. CYP inhibition profiling identified a second liability. **SN3-1** showed weak inhibition of most tested human CYP isoforms, with IC50 values greater than 10 μM. By contrast, it inhibited CYP2C19 with an IC_50_ of 1.59 μM (Figure 4D), raising concern for drug-drug interaction risk. Because CYP2C19 is involved in the metabolism of multiple clinical drugs, reducing this liability became a major optimization objective.

Metabolite analysis provided a structural explanation for these risks. High-resolution mass spectrometry indicated that the indane-containing side chain was a major region of phase I metabolism, consistent with oxidation around a benzylic position. The same region may also contribute to CYP2C19 binding because of its hydrophobic and aromatic character (Figure S4B). In parallel, the sulfonylurea core was associated with cleavage products containing an aniline-like fragment, consistent with a structural alert for drug-induced liver injury (Figure S4C). These observations converted the **SN3-1** optimization problem into two linked design tasks: block the metabolic soft spot around the indane region and replace or remodel the sulfonylurea-associated liability while preserving NLRP3 activity (Figure 4C).

#### 2.3.2. Iterative Structural Optimization and Discovery of B9

We next applied RCDO to SN3-1 through two iterative DMTA cycles (Figure 4E). The NLRP3 binding pocket and the three-dimensional conformation of SN3-1 provided the structural context for molecular generation. The reward system incorporated the same four-level logic used in the broader RCDO framework: topological validity, conformational plausibility, binding-mode complementarity and target-specific bioactivity. For the NLRP3 campaign, the target-specific reward was aligned with cellular NLRP3 inhibition and the experimentally defined ADME liabilities, especially CYP2C19 inhibition and microsomal stability (Table S6, and Figure S4D-E). Candidate molecules were then triaged through Pareto-frontier recommendation and expert inspection before synthesis and testing.

The first round focused on the metabolically vulnerable indane region. RCDO-guided generation prioritized substitutions expected to block oxidative metabolism while maintaining productive NLRP3 pocket occupancy. A focused set of Round I analogs was synthesized (**B1**-**B7**) and evaluated for inhibition of NLRP3 inflammasome activation. Among these, **B3** showed the strongest activity, with an IC_50_ of 2 nM, representing an approximately fourfold potency gain relative to **SN3-1** (Table S7). However, **B3** still retained the sulfonylurea warning motif which has been associated with development-limiting safety concerns. Accordingly, prior to the second round of molecular generation, we fixed the R_1_ substituent as an isopropyl group, which served as the core scaffold for the second round of design. The second round therefore shifted from local metabolic-site blocking to scaffold-level liability reduction. RCDO-guided scaffold hopping explored aminosulfonylurea analogs designed to reduce the electronic and cleavage liabilities of the original core while preserving three-dimensional similarity to the active scaffold (**B8**-**B13**). The best second-round compound was **B9**, which retained potent cellular activity with an IC_50_ of 10 nM (Table S7 and Figure 4G). The optimized compound **B9** showed a substantially improved developability profile (Figure 4H-J). In liver microsomes, **B9** was stable in both mouse and human systems, with high parent remaining after 60 minutes in mouse microsomes and no meaningful degradation (Figure 4H). CYP profiling showed that **B9** no longer strongly inhibited the tested major CYP isoforms and that its CYP2C19 IC_50_ shifted to 30 μM, an approximately 20-fold reduction in liability relative to **SN3-1** (Figure 4I-J). In *in vivo* pharmacokinetic studies in mice, oral administration of **B9** produced markedly elevated systemic drug exposure. The area under the concentration-time curve (AUC) was 5857 ng·mL⁻¹·h, and its oral bioavailability reached 96.4%. A dose-normalized comparison with **SN3-1** showed that **B9** exhibited significantly higher oral exposure (Table S8). Repeat-dose tolerability was evaluated in mice treated orally with **B9,** H&E staining of major organs did not show the clear liver injury pattern observed with SN3-1 (Figure 5A), and serum AST and ALT levels were not significantly elevated across the tested **B9** dose range (Figure 5B-C). These findings indicate that **B9** reduced **SN3-1**’s liver-liability signal under the tested repeat-dose conditions, confirming that **B9** is an optimized NLRP3 lead compound.

**Figure 5.**
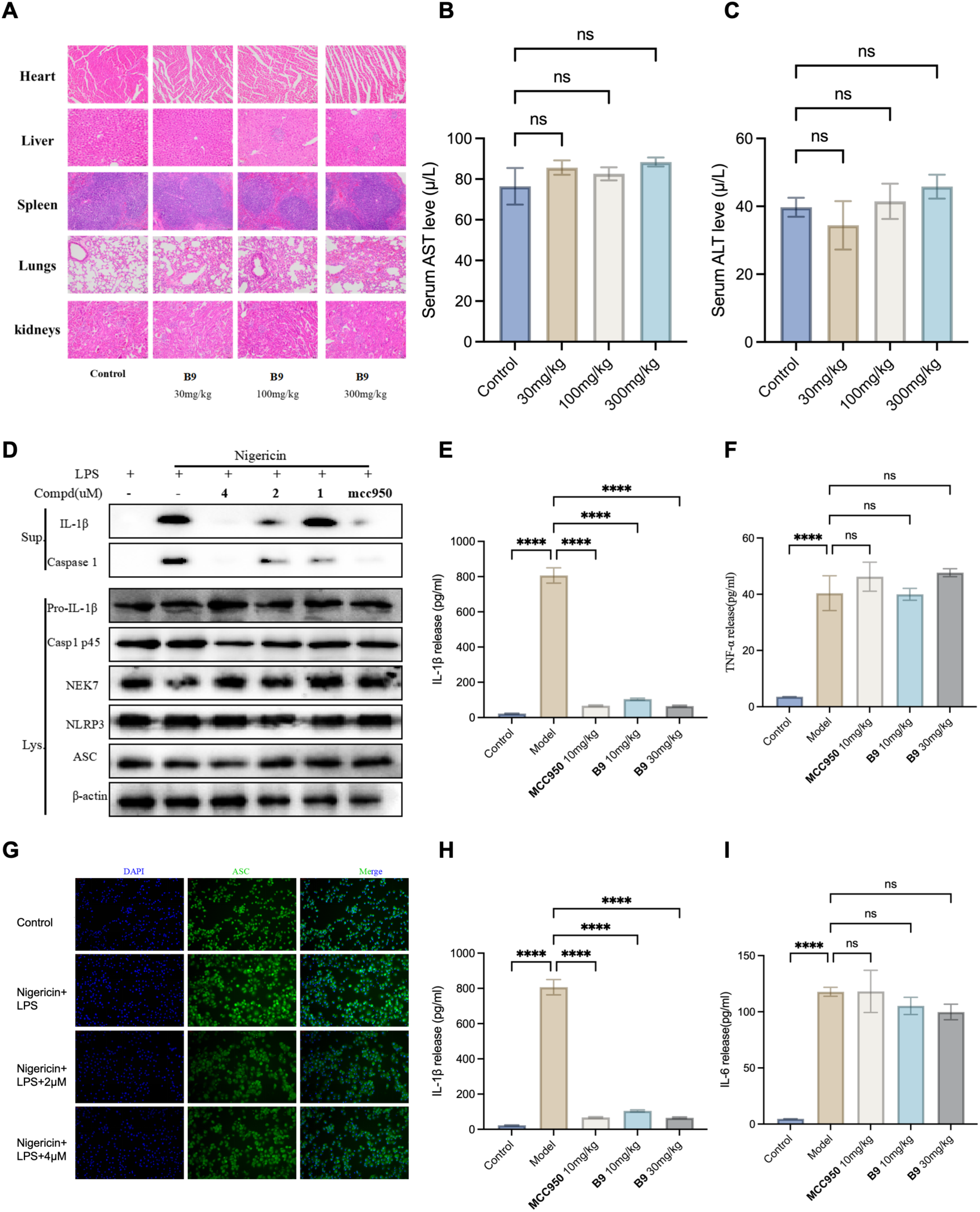
*In vitro* and *in vivo* activity and safety profile of Compound B9. **(A)** Representative H&E staining images of major organs collected after repeated oral administration of Compound **B9**, showing the histopathological safety profile of **B9** treatment. **(B)** Serum AST levels after repeated oral **B9** dosing (n = 3). (**C)** Serum ALT levels after repeated oral **B9** dosing (n = 3). **(D)** Immunoblot analysis of NLRP3 inflammasome activation following **B9** treatment, including inflammasome-associated signaling proteins and downstream inflammatory markers. **(E)** IL-1β levels in the LPS-induced acute inflammation model following treatment with **MCC950** or Compound **B9** (n = 3). **(F)** TNF-α levels in the LPS-induced acute inflammation model following treatment with **MCC950** or Compound **B9** (n = 3). (**G**) Representative immunofluorescence images of ASC speck formation, with DAPI-stained nuclei and ASC signals shown to visualize inflammasome assembly (n = 3). (**H**) IL-1β levels in the MSU-induced mouse peritonitis model following treatment with **MCC950** or Compound **B9** (n = 3). (**I**) IL-6 levels in the MSU-induced mouse peritonitis model following treatment with **MCC950** or Compound **B9** (n = 3). Data are presented as mean ± SEM where applicable. Statistical significance is indicated in the figure. Statistical significance was assessed using ordinary one-way ANOVA: ns (not significant), *p < 0.05, **p < 0.01, ***p < 0.001, ****p < 0.0001.

#### 2.3.3. *In Vitro* and *In Vivo* Activity Evaluation of B9

We next evaluated whether the improved ADME profile of **B9** preserved NLRP3-selective pharmacology. Immunoblot analysis showed that **B9** reduced mature IL-1β and cleaved caspase-1 after NLRP3 inflammasome activation, while basal pathway components, including NLRP3, ASC, pro-IL-1β and pro-caspase-1, were not broadly depleted (Figure 5D). This pattern indicates that **B9** did not simply suppress pathway protein expression, but interfered with inflammasome activation or assembly after priming.

The inhibitory activity of **B9** was evident against diverse NLRP3-activating stimuli, **B9** suppressed IL-1β maturation and caspase-1 activation in cells stimulated with nigericin, ATP or MSU. By contrast, **B9** exhibited negligible effects on inflammasome activation mediated by AIM2 or NLRC4 under the tested conditions (Figure S5A-B). These results confirm that **B9** selectively inhibits the NLRP3 inflammasome, rather than causing non-specific, broad suppression of all inflammasome cascades. ASC speck formation was then assessed as a cellular marker of NLRP3 inflammasome assembly. Confocal imaging showed that **B9** reduced ASC puncta in activated macrophages (Figure 5G). Because ASC oligomerization is a central assembly checkpoint downstream of NLRP3 activation, these data place the functional effect of **B9** at or before the ASC assembly step.

Finally, **B9** was tested in disease-relevant inflammatory models. In an LPS-induced systemic inflammation model, **B9** lowered IL-1β release while TNF-α was not broadly suppressed, supporting selective attenuation of NLRP3-dependent cytokine output rather than generalized immune suppression (Figure 5E-F). In an MSU-induced peritonitis model, **B9** reduced IL-1β in inflammatory readouts and had no effect on the level of IL-6 (Figure 5H-I). After intervention with **B9** at a dose of 30 mg/kg, the proportion of mature neutrophils in the mouse peritoneal lavage fluid decreased significantly to 1.17%, close to the baseline level of the control group. The above experimental results indicate that **B9** can effectively inhibit the recruitment and infiltration of key myeloid cells during acute inflammation, further demonstrating its favorable anti-inflammatory protective effect i*n vivo* (Figure S5C).

Together, following two rounds of molecular optimization with RCDO, we successfully addressed the hepatotoxic liability of our precursor Compound **SN3-1** and acquired Compound **B9**. While maintaining potent bioactivity, **B9** substantially alleviates potential liver toxicity and greatly enhances its druggability. Collectively, these results fully demonstrate that RCDO possesses the capacity for targeted optimization to attenuate the toxicity of small-molecule drugs.

### 2.4. Novel Scaffold Discovery and RCDO-Guided Optimization of NSD3 Inhibitors for Lung Squamous Cell Carcinoma

#### 2.4.1. Novel Scaffold Discovery of NSD3 Inhibitors Using Virtual Screening

Lung squamous cell carcinoma (LUSC) is a frequently recurring malignancy with few targeted treatment options and poor survival at advanced stages. Among its candidate drivers, nuclear receptor binding SET domain protein 3 (NSD3) has emerged as a direct oncogenic driver of this disease. NSD3 lies within the chromosome 8p11.2 amplicon, and its overexpression increases mono- and dimethylation of histone H3 at lysine 36 (H3K36me1/2). The resulting chromatin decompaction and transcriptional activation are propagated through BRD4 and c-MYC to drive proliferation, migration, and survival, and ultimately tumor progression (Figure 6A). Functional studies have established that NSD3 is required for LUSC initiation and progression, defining a high-value but as-yet-undrugged target. No NSD3 inhibitor has reached the clinic.

**Figure 6.**
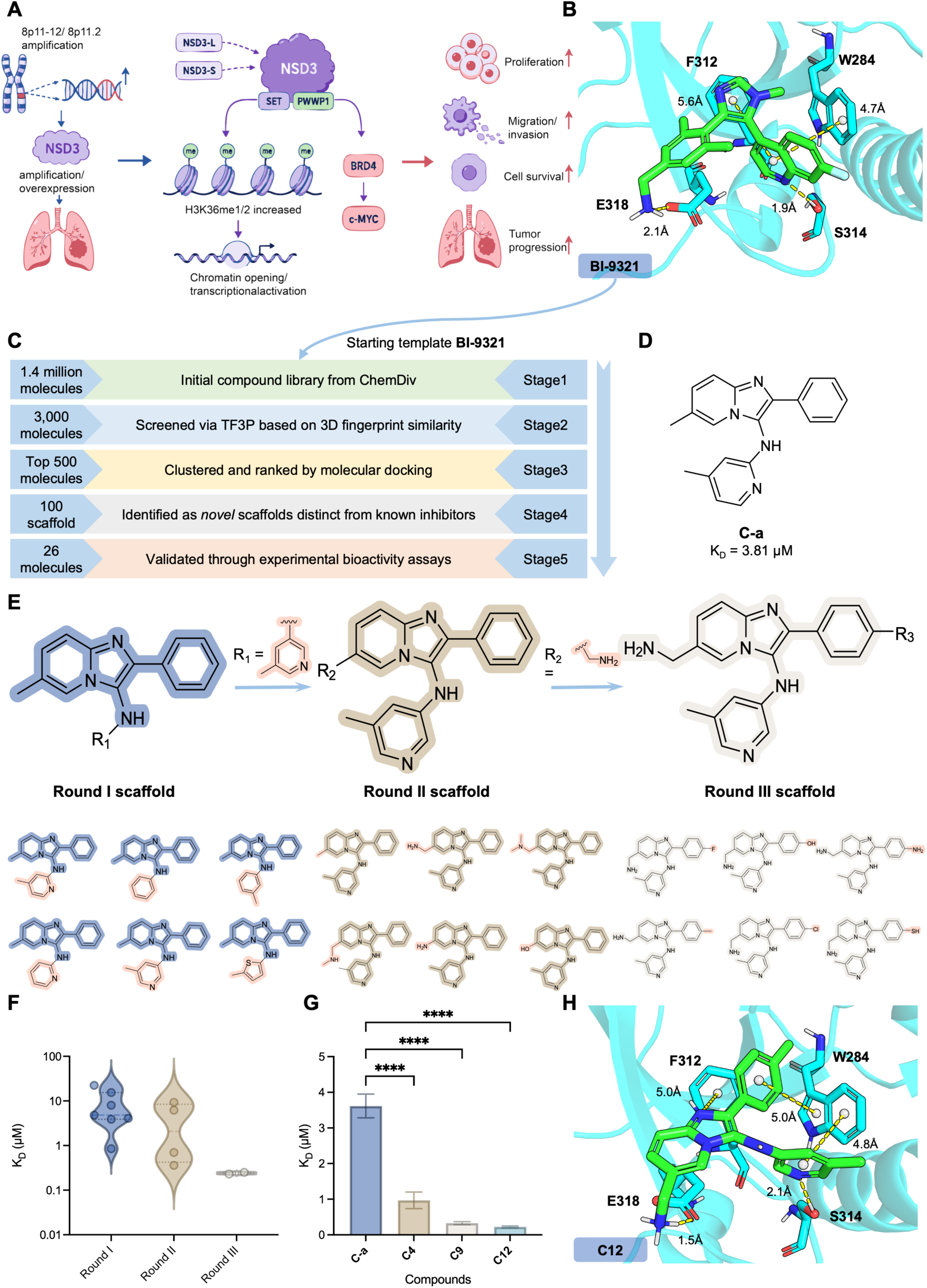
Discovery and RCDO-guided optimization of NSD3 PWWP1 inhibitors. **(A)** Schematic illustrating how NSD3 amplification or overexpression promotes tumor progression through H3K36me1/2-associated chromatin activation and BRD4–c-MYC signaling. **(B)** Binding pose of Compound **BI-9321** within NSD3 PWWP1 pocket, with surrounding residues and selected intermolecular distances indicated. **(C)** Virtual screening workflow based on Compound **BI-9321**. Starting from the ChemDiv library, successive filtering steps yielded 26 candidate compounds, followed by SPR validation identifying Compound **C-a**. **(D)** Chemical structure of screening hit **C-a** and its corresponding K_D_ value (3.81 μM). **(E)** Three-round scaffold-optimization strategy showing structural and substituent modifications used to generate Round I, Round II and Round III compound series. **(F)** Distribution of K_D_ values among compounds generated during each optimization round. **(G)** Stepwise comparison of K_D_ values for Compounds **C-a, C4**, **C9**, and **C12**, showing increased binding of optimized compounds relative to starting hit **C-a**. **(H)** Binding mode of optimized Compound **C12** within NSD3 PWWP1 binding pocket. Statistical significance was assessed using one-way ANOVA: ns (not significant), *p < 0.05, **p < 0.01, ***p < 0.001, ****p < 0.0001.

PWWP1 is a non-catalytic reader domain that anchors NSD3 to chromatin by engaging nucleosomal histone and DNA, thereby stabilizing substrate presentation during catalysis. Because the PWWP1 sequence diverges markedly among NSD family members, this domain is an attractive site for achieving selectivity for NSD3. The chemical probe **BI-9321** (Figure 6B) was recently identified as a selective antagonist of the NSD3 PWWP1 domain, and its co-crystal structure with the domain has been solved (PDB: 6G2O), providing a template for structure-based design of NSD3 inhibitors.

We therefore set out to identify a novel NSD3 scaffold through a virtual screening campaign (Figure 6C). Using **BI-9321** as a 3D reference geometry, we screened against the NSD3 PWWP1 binding pocket. A 1.4-million-compound ChemDiv library was first filtered with the TF3P^19^ fingerprint, a deep capsule-network descriptor that encodes electrostatic and van der Waals shape, yielding 3,000 candidates that matched the 3D profile of **BI-9321**. TF3P ranking retained the top 500, Bemis–Murcko scaffold clustering condensed these into 100 scaffold classes, and structure-based docking against the PWWP1 pocket, followed by manual inspection, nominated 26 compounds for purchase (Table S9). Surface plasmon resonance against recombinant NSD3 PWWP1 confirmed binding of the top hit, Compound **C-a** (K_D_ = 3.81 μM) (Figure 6D). Compound **C-a** defined a scaffold distinct from that of BI-9321 and served as the lead for the subsequent closed-loop optimization.

#### 2.4.2. Iterative Structural Optimization and Discovery of Compound C12

Compound **C-a** only exerted weak NSD3 binding affinity at the micromolar level. To further improve its target binding affinity, we applied RCDO to drive scaffold evolution across three iterative optimization rounds (Figure 6E). The reward system was configured to retain the core pharmacophore of **C-a** while implementing sequential generative diversification at the R_1_, R_2_, and R_3_ substituent positions, with multi-level rewards (Table S10, and Figure S6B-D).

In the first optimization round, we fixed all other scaffold moieties to systematically explore substitutions at the R1 position, followed by chemical synthesis of analogs **C1–C7** (Table S11). Among these candidates, Compound **C4** exhibited favorable NSD3 binding affinity with a K_D_ value of 0.85 μM. Subsequently, we locked the R_1_ substituent as the moiety derived from C4 and performed structural diversification at the R_2_ site. Meanwhile, all binding activity data acquired from the first-cycle compounds were integrated into the reward system to drive second-round molecular generation, from which we synthesized Compounds **C8-C11**. Compound **C9** stood out with improved NSD3 binding affinity (K_D_ = 0.33 μM, Figure S6A). On this basis, we launched the third round of molecular generation and synthesized **C12** and **C13**. Compound **C12** achieved outstanding NSD3 PWWP1 binding affinity, with a K_D_ of 0.22 μM (Figure S6A, and Table S11). **C12** maintains key interactions with residues F312, W284, E318, and S314, showing an improved binding conformation relative to the initial template (Figure 6H).

The iterative optimization of NSD3-targeting compounds focused primarily on enhancing target binding affinity. As illustrated in Figure 6F, the NSD3 binding potency of synthetic compounds progressively improved across the three cycles, and the representative lead from each round (**C4**, **C9** and **C12**) displayed stepwise elevated binding affinity (Figure 6G, and Figure S6A). The final optimized compound **C12** delivered an approximately 18-fold improvement in NSD3 binding affinity compared with the starting compound **C-a**. Consistent with the above lead discovery campaign, the RCDO framework demonstrated outstanding performance in boosting NSD3 binding potency. This framework narrows the molecular search space stepwise via experimentally validated substituent constraints, and leverages binding data from each cycle to refine the sampling strategy for subsequent rounds of molecular generation.

## 3. Discussion

Drug discovery has historically relied on an DMTA process, in which experimental outcomes continuously reshape medicinal chemistry decisions. However, most AI-driven molecular design approaches remain largely open-loop: computational models generate candidate molecules according to predefined objectives, while experimental results are mainly used for downstream evaluation. Here, RCDO demonstrates that experimental feedback can be transformed from a downstream evaluation step into a driving force for molecular evolution. By coupling a 3D molecular generative model UniLingo3DMol^7^ with a multi-level reward system updated after DMTA design cycle using experimental measurements from all synthesized compounds, including inactive or developability-failed compounds, RCDO continuously aligns generative exploration with accumulated wet-lab data and adapts design strategies across successive DMTA cycles.

RCDO extends a broader emerging concept of feedback-driven AI in biological discovery. Recent studies such as EVOLVEpro^20^ have demonstrated experimental feedback-driven evolution in protein sequence space. However, applying this principle to small molecule optimization presents a distinct challenge because drug-like molecules must simultaneously satisfy constraints involving 3D binding geometry, chemical feasibility, pharmacokinetic properties, metabolism, and safety. Rather than only prioritizing existing candidates, RCDO enables experimental measurements to directly reshape the molecular generation model, extending closed-loop learning from biological sequences and compound prioritization to the evolution of 3D small molecule structures.

This study also illustrates that molecular optimization in practice requires navigation through a multi-dimensional property landscape rather than maximization of a single objective. The ROR1 and NLRP3 campaigns illustrate this challenge: improving target engagement alone is insufficient when permeability, metabolic stability, CYP inhibition, or toxicity liabilities limit therapeutic potential. By incorporating experimentally measured properties into subsequent DMTA cycles, RCDO progressively shifted molecular populations toward more balanced solutions.

Several limitations remain. First, the retrospective benchmark scores structural rediscovery of validated analogs rather than direct biological activity, as high-activity candidates are defined by ECFP Tanimoto similarity above 0.9 to a known active analog. This proxy may fail near activity cliffs and structurally penalizes active molecules distinct from the historical series. Second, RCDO remains a human-in-the-loop (HITL) framework, as medicinal chemistry expertise is still required for evaluating synthetic feasibility, structural novelty, and strategic prioritization. However, this involvement reflects a broader challenge in AI-driven discovery: many aspects of medicinal chemistry represent implicit expert knowledge that has not yet been fully encoded computationally. Future systems may reduce this dependency by integrating reaction-aware generation, automated synthesis, richer experimental representations, and improved modeling of biological mechanisms and uncertainty. Nevertheless, the goal may not be complete replacement of medicinal chemists, but rather the development of collaborative systems in which computation expands chemical exploration while humans provide mechanistic insight and strategic direction.

More broadly, RCDO suggests a transition in AI-assisted drug discovery from static generation into continuous experimental adaptation. The future of computational chemistry may depend not only on generating more molecules, but on developing models that learn from experimental interactions and progressively improve their own discovery strategies. By demonstrating that experimental feedback can drive the evolution of 3D small molecules, RCDO provides a framework for integrating computation and experimentation into a unified learning process for drug discovery.

## 4. Methods

### 4.1. Base Model for Molecular Directed Optimization

The basic model of RCDO framework is our previously developed UniLingo3DMol, a transformer-based encoder-decoder model for conditional 3D molecular generation. We selected UniLingo3DMol for its native support for diverse generation scenarios, which lets our optimization workflow adapt to flexible starting points: UniLingo3DMol supports both *de novo* molecular generation, where we explore chemical space from scratch, and fragment-retained generation, where we refine existing core scaffolds while preserving their critical binding interactions.

UniLingo3DMol provides a generalizable initialization for our RL framework, removing the need to train a molecular generative model from scratch and allowing the directed optimization to converge quickly toward high-quality molecules. The full architectural details of UniLingo3DMol, including its DSMILES-based representation syntax and multi-head generation mechanism, are given in our previous study^7^.

### 4.2. Hierarchical Multi-level Reward System

To enable efficient molecular directed optimization, we developed a systematic, multi-level reward system organized into four hierarchical levels: (a) topological validity, (b) conformational plausibility, (c) binding mode complementarity, and (d) target-specific bioactivity. This architecture encapsulates core drug design constraints, guiding the optimization process toward high-quality chemical space.

#### Topological Level

The topological level forms the foundation of our reward system, addressing the core requirement of generating chemically valid, drug-like, synthetically accessible molecules. By enforcing strict medicinal chemistry constraints, this level filters out chemically implausible molecular structures before higher-level optimization. Without robust 2D topology, improvements in binding affinity are computationally meaningless. Topological rewards thus confine directed optimization to a chemically navigable, realistically synthesizable chemical space. We implemented several rewards at this level to constrain synthesizability, molecular flexibility, functional group compatibility, and physicochemical properties that can be calculated from the molecular topological structure. More details of topological level rewards are provided in Section S1.1.

#### Conformational Level

Building on the topological level, the conformational level assesses whether a molecule’s 3D geometry is physically plausible and energetically accessible. Although UniLingo3DMol is able to generate molecules with effective connectivity, these molecules may contain structural defects such as spatial clashes or unrealistic torsion angles. By penalizing the high-energy states of molecules, our reward system ensures molecules can avoid unreasonable conformations. Here, we implemented rewards to evaluate internal atomic strain and protein-ligand binding energetics, with full details provided in Section S1.2.

#### Binding Mode Level

Next, the binding mode level rewards evaluate complementarity between the ligand and protein pocket, shifting focus from the isolated molecule to the protein-ligand complex. This level rewards specific noncovalent interaction (NCI) pattern, such as hydrogen bonding networks and hydrophobic shielding, that are critical for molecular recognition. The NCI reward quantifies how well the ligand exploits available interaction partners in the binding pocket (e.g., hydrogen bonds, salt bridges, π-π stacking), ensuring geometric and electrostatic complementarity rather than simple shape matching, with details provided in Section S1.3.

#### Target-specific Bioactivity Level

Finally, the target-specific bioactivity level forms the apex of the reward system, incorporating quantitative structure-activity relationship (QSAR) predictions to estimate bioactivity. For example, cellular activity, biochemical activity, etc. While preceding levels ensure a molecule is capable of binding, the bioactivity reward directs optimization toward genuine pharmacological efficacy. Critically, this level bridges dry-lab computational simulations and wet-lab experimental validation: by constructing QSAR models based on empirical biological activity data, it ensures generated molecules are not merely theoretical ideals, but biologically relevant leads ready for experimental testing. We trained task-specific QSAR models using corresponding wet-lab experimental data, with full details provided in Section S1.4.

### 4.3. Multi-level Rewards Aggregation Strategy

To distill multi-level rewards into a unified scalar objective for RL policy updates, we designed a hierarchical aggregation strategy tailored to the distinct functional roles of each reward component. Unlike standard weighted summations that treat all objectives equally, our approach explicitly decouples hard constraints from soft optimization targets. We route each reward through one of two aggregation operators, including multiplicative and additive operators.

The multiplicative operator enforces non-negotiable physical constraints, such as chemical validity and functional group compatibility, by acting as a strict veto mechanism. A zero score from any multiplicative constraint collapses the total reward to zero, ensuring the RL policy receives no positive gradient for infeasible molecules and thereby conserving model capacity for viable chemical space. Conversely, the additive operator governs soft constraints, representing tunable properties like binding affinity or bioactivity. These components contribute a weighted score, establishing a continuous gradient landscape that actively steers the policy toward high-performing structural candidates. Formally, for a set of *N* reward components, the aggregated reward *R* is defined as:

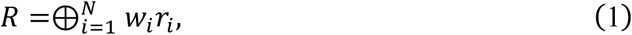

where *w_i_* and *r_i_* denote the weight and score of the *i*-th reward component, respectively. The operator ⊕ indicates the operator-specific accumulation, which is initialized at zero for additive and at one for multiplicative operators, and *R* provides the reward signal for our RL learning policy.

### 4.4. Reinforcement Learning Empowered Molecular Directed Optimization

We formulate molecular directed optimization as a RL problem, wherein the pre-trained UniLingo3DMol acts as the generative policy *π_θ_*, guided by the aggregated scalar reward *R*. To update the policy network, we employ the group relative policy optimization (GRPO) strategy. By leveraging intra-group statistics to estimate the advantage, GRPO circumvents the computational overhead and memory footprint associated with training a separate value network.

#### 4.4.1. Group Normalization and Advantage Estimation

Rather than relying on a parameterized value function to establish a baseline, GRPO evaluates candidate molecules relatively. During each RL epoch, for a given binding pocket, the policy autoregressively samples a group of *G* distinct candidate molecules. We evaluate the aggregated reward *R_i_* for each molecule within this group. The advantage *A_i_* for the *i*-th molecule is then computed by standardizing its reward against the empirical distribution of the group:

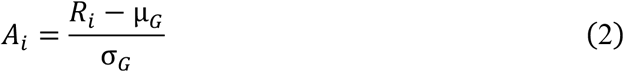

where µ*_G_* and σ*_G_* are the mean and standard deviation of the rewards across the *G* sampled molecules.

#### 4.4.2. Policy Optimization Objective

Driven by these group-normalized advantages, the policy is updated by minimizing a clipped surrogate loss, augmented with a Kullback-Leibler (KL) divergence penalty to ensure training stability:

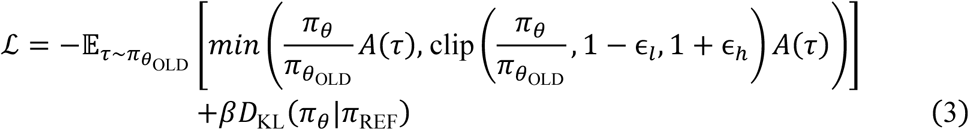

where *A*(*τ*) denotes the group-normalized advantage, *π*_*θ*_OLD__ is the policy prior to the update step, *π*_REF_ is the frozen pre-trained reference model, *β* controls the strength of the KL penalty, and *D*_KL_ denotes the KL divergence. In practice, we set ɛ*_l_* = ɛ*_h_* = 0.2 and *β* = 0.05. To compute this objective, we calculate the log-probabilities for each generated molecule under both the active policy *π_θ_* and the reference policy *π*_REF_. The exponential difference in log-probabilities between the active and old policies defines the policy ratio *π_θ_*⁄*π*_*θ*_OLD__. Concurrently, the logits from the reference model supply the baseline for the KL divergence penalty *D*_KL_(*π_θ_*|*π*_REF_). This penalty acts as a vital regularization mechanism, anchoring the optimized model to the pre-trained chemical distribution and preventing divergence when optimization is guided by noisy reward signals.

#### 4.4.3. Progressive Multi-Round Optimization Strategy

To align with the empirical DMTA cycle intrinsic to drug discovery, our directed optimization operates as a progressive, HITL iterative pipeline rather than a single-shot generative task. The hyperparameter settings of RCDO are shown in Table S12.

##### Iteration of Scaffold

As the optimization progresses across multiple rounds, the generative degrees of freedom are systematically constrained. Initially, the RL policy explores a broad combinatorial space across multiple substitution sites on a parent molecular core. When early round wet-lab experiments validate the efficacy of specific structural motifs, these optimal R-groups are explicitly locked into the input prompt for subsequent generations. This progressive fixation mechanism continuously narrows the explorable chemical space, transitioning the model from broad structural elaboration to highly targeted local refinement of the remaining undetermined topological regions.

##### Iteration of Rewards

Concurrently, the multi-dimensional reward landscape evolves by encoding experimental feedback. After each generation step, recommended candidate molecules are triaged via expert visual inspection and advanced to wet-lab synthesis and validation. The newly acquired experimental data are directly utilized to retrain and calibrate predictive scoring functions, such as QSAR models, thereby continuously enhancing their predictive fidelity. Furthermore, structural insights extracted from these experimental cycles, such as the identification of unexpected unstable motifs or metabolic liabilities, are formalized and integrated into the reward framework as novel functional group alerts. Consequently, the RL environment becomes progressively more stringent and accurately aligned with the physical realities of the drug discovery project.

#### 4.4.4. Pareto-Frontier Molecule Recommendation strategy

The final stage of each optimization round selects which generated molecules to advance to expert inspection and wet-lab validation. A naive strategy that retains only molecules with the highest aggregated reward tends to over-concentrate on a single high-scoring region of chemical space and to discard candidates that excel on individual objectives but are penalized on others. Conversely, forwarding the entire generated library overwhelms downstream triage and dilutes experimental throughput. To balance these extremes, we adopt a Pareto-frontier recommendation strategy operating over the individual reward dimensions, such as topological favorability, conformational stability, binding-mode complementarity, and predicted bioactivity. Rather than collapsing these objectives into a single scalar for ranking, we retain every molecule that is non-dominated, that is, optimal on at least one reward dimension without being strictly inferior on all others, thereby assembling a compact molecular set that spans the trade-off surface among competing pharmacological objectives. This Pareto-based set preserves candidates that represent the best achievable value for each individual property, maximizing the diversity and information content of each experimental cycle while ensuring that no single optimization objective is silently sacrificed during selection.

### 4.5. Biological Assay Methods

All animal experiments were performed following the protocols evaluated and approved by the Animal Ethics Committee of Peking University (Ethics Approval Number: DLASBE0111)

#### 4.5.1. Chemical Synthesis and Compound Characterization

Recommended compounds were selected from each Pareto set after medicinal-chemistry inspection and synthesized according to target-specific synthetic routes. Final compounds and key intermediates were characterized by nuclear magnetic resonance spectroscopy and high-resolution mass spectrometry before biological testing.

#### 4.5.2. Surface Plasmon Resonance Binding Assays

SPR assays were used to quantify direct binding of optimized compounds to ROR1, NLRP3 or NSD3 PWWP1. For ROR1 and NSD3, proteins were diluted in pH 4.0 sodium acetate buffer to 100 ng/μL and immobilized on CM5 sensor chips by amine coupling to an approximate immobilization level of 15,000 response units. Reference flow cells were activated and blocked without protein immobilization. PBS-P was used as running buffer. Compound dilution series were prepared with 5% DMSO, and 4.5-5.8% DMSO calibration solutions were used to correct bulk solvent effects. Compounds were injected sequentially over the immobilized and reference surfaces, and sensorgrams were recorded through association and dissociation phases. The chip surface was regenerated by buffer wash after each injection.

For NLRP3 binding assays, NLRP3 protein was diluted to 20 μg/mL and immobilized on a CMS sensor chip to approximately 5,000-10,000 response units by standard amine coupling. Residual activated groups were blocked with 1 M ethanolamine-HCl. Small-molecule analytes were serially diluted in running buffer containing 10 mM HEPES and 150 mM NaCl at pH 7.4 and injected at 30 μL/min. Raw signals were corrected by double referencing against the reference flow cell and blank buffer injections. Corrected sensorgrams were globally fit to a 1:1 binding model to calculate association rate, dissociation rate and equilibrium dissociation constant.

#### 4.5.3. Cell Culture and Antiproliferative Assays

MDA-MB-231 cells were cultured in DMEM supplemented with 10% fetal bovine serum and penicillin-streptomycin at 37℃ with 5% CO_2_. ROR1-series antiproliferative activity was measured by MTT assay. Log-phase cells were seeded in 96-well plates at 5,000 cells per well, allowed to attach overnight, treated with serial concentrations of test compounds for 72 h, incubated with 5 mg/mL MTT for 4 h and dissolved in 150 μL dimethyl sulfoxide. Absorbance was measured at 490 nm, and IC_50_ values were calculated in GraphPad Prism 8 using a three-parameter concentration-response model.

#### 4.5.4. NLRP3 Inflammasome Cellular Assays

THP-1 monocytes were adjusted to 1*10^6^ cells/mL and differentiated with 50 ng/mL phorbol 12-myristate 13-acetate for 24 h. After washing and overnight recovery in RPMI 1640 containing 1% fetal bovine serum, cells were primed with 50 ng/mL lipopolysaccharide (LPS) for 3 h. Test compounds or 0.1% DMSO vehicle were added for 30 min before inflammasome activation. NLRP3 activation was induced with 200 μg/mL monosodium urate (MSU) crystals for 4 h, or with 5 mM ATP or 10 μM nigericin for 30 min. Bone-marrow-derived macrophages (BMDMs) were generated from C57BL/6 mouse femur and tibia marrow cells by 7-day differentiation in DMEM containing 10% fetal bovine serum, 30% L929 conditioned medium and antibiotics. BMDMs were seeded at 5 x 10^5 cells/mL and primed with 200 ng/mL LPS for 3 h before compound treatment and inflammasome activation.

To evaluate inflammasome selectivity, AIM2 and NLRC4 activation models were run in parallel. After LPS priming, AIM2 activation was induced by Lipofectamine 2000-mediated transfection of 1 μg/mL poly(dA:dT) for 4 h, and NLRC4 activation was induced by exposure to 100 ng/mL Salmonella typhimurium flagellin for 2 h. Cytokines in culture supernatants were quantified with commercial ELISA kits according to the manufacturers’ protocols.

#### 4.5.5. Immunoblotting, ASC Speck Imaging and Functional Cell Assays

For immunoblotting, treated cells were lysed in ice-cold RIPA buffer containing protease and phosphatase inhibitors. Lysates were cleared by centrifugation, protein concentrations were determined by BCA assay, and samples were separated by SDS-PAGE before transfer to PVDF membranes. Membranes were blocked, incubated with primary antibodies overnight at 4℃, washed with TBST, incubated with horseradish peroxidase-conjugated secondary antibodies and visualized by enhanced chemiluminescence. For ROR1 studies, antibodies included ROR1, phosphorylated ROR1, cleaved PARP, Bax, Bcl-2 and GAPDH. For NLRP3 studies, intracellular lysates and methanol/chloroform-precipitated supernatant proteins were used to assess inflammasome pathway components and activation products, including mature IL-1β and cleaved caspase-1.

ASC speck formation was assessed in PMA-differentiated THP-1 macrophage-like cells after inflammasome activation. Cells were fixed with 4% paraformaldehyde for 15 min, permeabilized with 0.1% Triton X-100 for 10 min, blocked with 5% bovine serum albumin for 1 h, incubated with anti-ASC antibody at 1:1,000 overnight at 4℃, washed, incubated with fluorophore-conjugated secondary antibody at 1:1,000 for 1 h and counterstained with DAPI. ASC puncta were imaged by confocal microscopy.

Colony-formation assays were used to evaluate long-term proliferative capacity. For ROR1 studies, MDA-MB-231 cells were seeded at 800 cells per well in six-well plates, treated with compound or DMSO after 24 h and cultured for 2 weeks. Colonies were fixed with 4% paraformaldehyde, stained with 0.01% crystal violet.

Scratch-wound migration assays were performed by scratching confluent monolayers with a 20 μL pipette tip, washing with PBS, replacing medium with serum-free medium and imaging wound closure over 0-48 hours.

#### 4.5.6. Kinase Selectivity Profiling

The kinase selectivity profile of the optimized ROR1 lead was evaluated with a broad kinase panel provided by Eurofins. Compound **A11** was tested at 0.5 μM across 360 kinase assays, and kinases showing measurable interaction under the tested condition were mapped in the selectivity profile.

#### 4.5.7. Microsomal Stability, PAMPA Permeability and CYP Inhibition

Human liver microsomal stability assays were used to evaluate metabolic turnover. Test compounds were incubated with human liver microsomes, NADPH and UDPGA at 37℃. At predefined time points, aliquots were quenched with acetonitrile, mixed, stored at -80℃ and centrifuged before LC-MS/MS analysis of remaining parent compound. Half-life was calculated as T_1/2_ = -0.693/k, where k was obtained from the slope of the linear fit between incubation time and the natural logarithm of the remaining parent fraction.

Parallel artificial membrane permeability assay (PAMPA) was used to evaluate passive membrane permeability of ROR1 analogs. Each filter membrane received 5 μL lecithin solution. Donor wells contained 300 μL of 10 μM test or control compound, and acceptor wells contained 300 μL PBS with 1% DMSO. Plates were incubated at approximately 60 rpm and 37℃ for 16 h. After incubation, 50 μL samples from acceptor, donor and blank solutions were mixed with 200 μL acetonitrile containing internal standard, centrifuged at 4℃ and 4,200 g for 15 min, diluted with water and analyzed by LC-MS/MS.

CYP inhibition assays were conducted in human liver microsome incubation systems to assess inhibition of major CYP450 isoforms. For NLRP3 compounds, CYP1A2, CYP2C9, CYP2C19, CYP2D6 and CYP3A4 were tested with isoform-specific probe substrates, including phenacetin, diclofenac, S-mephenytoin, dextromethorphan, midazolam and testosterone. Selective inhibitors were run as positive controls. Human liver microsome concentration was 0.5 mg/mL for CYP2C19 and 0.1 mg/mL for the other isoforms. Reaction mixtures were preincubated at 37℃ for 5 min and initiated with prewarmed NADPH. Incubation times were 5 min for CYP3A4, 10 min for CYP1A2 and CYP2C9, 20 min for CYP2D6 and 45 min for CYP2C19. Reactions were quenched with acetonitrile/methanol (1:1, v/v) containing tolbutamide as internal standard, and supernatants were analyzed by LC-MS/MS to calculate IC_50_ values.

#### 4.5.8. Metabolite Identification and Pharmacokinetics

For metabolite identification in the NLRP3 campaign, test compounds were incubated with primary human hepatocytes at 1.0*10^6^ cells/mL in William’s E medium at 37℃ with 5% CO_2_. Samples were quenched with acetonitrile at 0 or 120 min, centrifuged, dried under nitrogen, reconstituted in acetonitrile/water and analyzed by LC-UV-HRMS on an Orbitrap system. Data were processed with Compound Discoverer to assign metabolite structures.

Pharmacokinetic studies were performed in male ICR mice. For ROR1 compounds, oral dosing was conducted at 10 mg/kg, and intravenous dosing was performed at 1 mg/kg where absolute bioavailability was required. Blood samples were collected after oral dosing at 0.25, 0.5, 1, 2, 4, 6 and 8 h, and after intravenous dosing at 0.083, 0.25, 0.5, 1, 2, 4 and 8 h. Plasma was isolated by centrifugation at 4℃ and stored at -80℃ before LC-MS/MS analysis. For NLRP3 compounds, mice were fasted for 12 h and dosed intravenously at 2 mg/kg or orally at 10 mg/kg in 5% DMSO plus 10% cyclodextrin aqueous vehicle. Plasma samples were collected at 0.083, 0.25, 0.5, 1, 2, 4 and 8 h, processed by protein precipitation and quantified by LC-MS/MS in multiple reaction monitoring mode. Pharmacokinetic parameters were calculated by non-compartmental analysis.

#### 4.5.9. ROR1 *In Vivo* Efficacy and Safety Studies

All animal procedures were approved by the relevant institutional animal ethics committee and conducted according to approved protocols. For subcutaneous TNBC xenografts, log-phase MDA-MB-231 cells were washed with PBS and suspended in a 1:1 mixture of PBS and Matrigel at 5*10^7^ cells/mL. Each BALB/c nude mouse received 0.1 mL suspension containing 5*10^6^ cells by subcutaneous injection. When tumor volume reached approximately 50-100 mm^3^, mice were randomized into vehicle, Compound **59** and optimized-lead groups. For the final ROR1 lead experiment corresponding to Figure 3E, compounds were administered by oral gavage at 30 mg/kg every 3 days for 27 days. Tumor volume and body weight were measured at regular intervals. Tumor volume was calculated as length*width^2^*0.5. At endpoint, mice were euthanized, tumors were excised and weighed, and major organs were collected for histopathology.

For orthotopic TNBC studies, 3*10^6^ luciferase-expressing MDA-MB-231 cells were injected into the mammary fat pad of BALB/c nude mice. When tumors reached approximately 80-100 mm^3^, mice were randomized into vehicle and optimized-lead groups. The optimized lead was administered orally at 20 mg/kg every 3 days. In vivo bioluminescence imaging was performed after intraperitoneal injection of 100 μL D-luciferin, with images acquired approximately 10 min after substrate injection. Imaging was performed on days 7 and 43 after treatment initiation for the orthotopic model. At endpoint, mice were euthanized and major organs were collected for ex vivo fluorescence imaging and histopathological evaluation.

Acute toxicity was evaluated in ICR mice after a single oral dose of 1,000 mg/kg test compound or vehicle. Mice were monitored for 14 days for survival, clinical signs and body-weight changes. At the end of the observation period, major organs were collected for hematoxylin and eosin (H&E) staining.

#### 4.5.10. NLRP3 *In Vivo* Safety and Inflammatory Disease Models

Repeat-dose tolerability of NLRP3 compounds was evaluated in 8-10-week-old C57BL/6 mice. Mice were randomized into control and dose groups and treated orally for 12 consecutive days at 30-300 mg/kg. On the day after the final dose, peripheral blood was collected for liver and kidney chemistry assays, and heart, liver, spleen, lung and kidney tissues were collected for histopathological analysis.

For the LPS-induced systemic inflammation model, 8-week-old C57BL/6 mice were randomized to treatment groups. Mice received test compound at 10 or 30 mg/kg, or positive-control inhibitor at 10 mg/kg, 30 min before intraperitoneal injection of 20 mg/kg LPS; control animals received PBS. Six hours after LPS challenge, serum was collected and IL-1β and other cytokines, including TNF-α where indicated, were measured by ELISA. Major organs were collected for H&E staining.

For the MSU-induced peritonitis model, 8-week-old C57BL/6 mice were randomized to treatment groups. Treatment groups received intraperitoneal injections of test compound for 3 consecutive days, and control mice received an equal volume of PBS. Thirty minutes after the final pretreatment, acute peritonitis was induced by intraperitoneal injection of 1 mg MSU crystals in 200 μL PBS. Six hours later, mice were euthanized and peritoneal cavities were lavaged with 5 mL PBS. IL-1β and IL-6 in lavage fluid or serum were quantified by ELISA as indicated in the figure legends, and neutrophil infiltration in peritoneal lavage fluid was analyzed by flow cytometry.

#### 4.5.11. NSD3 PWWP1 Protein Preparation

NSD3-PWWP1 (amino acids 247-398) was expressed and purified for SPR-based binding assays using an ampicillin-resistant bacterial glycerol stock encoding a His/GST-tagged NSD3-PWWP1 fusion protein with an rTEV cleavage site. The glycerol stock was plated on LB agar containing 100 μg/mL ampicillin and incubated overnight at 37°C. A single colony was inoculated into LB medium containing 100 μg/mL ampicillin and grown at 37°C with shaking at 200 rpm until the OD600 reached 0.6-0.8. Protein expression was induced with 0.5 mM IPTG for 16 h at 15°C. Cells were harvested at 4°C by centrifugation at 16,000 rpm, resuspended and lysed by sonication for 30 min. The lysate was clarified by centrifugation at 12,000 rpm for 20 min, and the supernatant was loaded onto Ni-NTA agarose resin. Bound His/GST-NSD3-PWWP1 was eluted using imidazole-containing buffer and analyzed by SDS-PAGE followed by Coomassie blue staining. The GST tag was then removed by rTEV protease digestion according to the manufacturer’s instructions. The digested sample was further purified using GST agarose resin to remove GST-containing species and uncleaved fusion protein. Fractions containing cleaved NSD3-PWWP1 were collected, yielding NSD3-PWWP1 protein with >90% purity.

#### 4.5.12. Histology and Statistical Analysis

Tissues were fixed in 4% paraformaldehyde or 10% formalin, dehydrated, embedded in paraffin, sectioned and stained with H&E. Slides were imaged by light microscopy. For NLRP3 safety and inflammatory models, tissue lesions were graded semi-quantitatively on a 0-4 scale, where 0 indicated no obvious lesion and 1-4 indicated minimal, mild, moderate and severe pathology, respectively.

Data are presented as mean ± standard error of the mean as indicated in the corresponding figure legends. IC_50_ values were calculated from concentration-response curves. Statistical comparisons in the main figures were performed with two-sided Student’s t-tests for two-group comparisons and with ordinary one-way ANOVA for multi-group comparisons where indicated; significance thresholds were denoted as ns, not significant, *P < 0.05, **P < 0.01, ***P < 0.001 and ****P < 0.0001.

## Code and Data Availability

The RCDO source code will be released publicly following acceptance of this manuscript. All experimental data from the DMTA procedure are available in the Supplementary Tables.

## Acknowledgements

This work was supported by China Postdoctoral Science Foundation (No. 2025M781450), and National Natural Science Foundation of China (No. 82504563).

## Author Contributions

## References

1. Guan, J., et al. 3D equivariant diffusion for target-aware molecule generation and affinity prediction. in International Conference on Learning Representations (2022).

2. Feng, W. et al. Generation of 3D molecules in pockets via a language model. *Nat*. Mach. Intell. 6, 62–73 (2024).

3. Qu, Y., et al. MolCRAFT: Structure-based drug design in continuous parameter space. in International Conference on Machine Learning (2024).

4. Chen, S. et al. Deep lead optimization enveloped in protein pocket and its application in designing potent and selective ligands targeting LTK protein. Nat Mach Intell 7, 448–458 (2025).

5. Loeffler, H. H. et al. Reinvent 4: Modern AI-driven generative molecule design. J. Cheminf. 16, 20 (2024).

6. Zhang, X. et al. Steering semi-flexible molecular diffusion model for structure-based drug design with reinforcement learning. Sci. Adv. 12, eady9955 (2026).

7. Huang, B., et al. A unified language model bridging de novo and fragment-based 3D molecule design. Preprint at 10.21203/rs.3.rs-8558464/v1 (2026).

8. Yuan, K. et al. Discovery of potent, selective, and orally bioavailable DYRK2 inhibitors for the treatment of prostate cancer. J. Med. Chem. 66, 16235–16256 (2023).

9. Dong, G. et al. Discovery and evaluation of DA-302168S as an efficacious oral small-molecule glucagon-like peptide-1 receptor agonist. J. Med. Chem. 68, 9555–9583 (2025).

10. Cheng, X. et al. Discovery of oxime ether derivatives as second-generation PRMT5 inhibitors for the treatment of triple negative breast cancer. J. Med. Chem. 69, 2647–2665 (2026).

11. Santos-Martins, D. et al. Accelerating AutoDock4 with GPUs and gradient-based local search. J. Chem. Theory Comput. 17, 1060–1073 (2021).

12. Maqbool, M., Bekele, F. & Fekadu, G. Treatment strategies against triple-negative breast cancer: An updated review. BCTT **Volume** 14, 15–24 (2022).

13. Bray, F. et al. Global cancer statistics 2022: GLOBOCAN estimates of incidence and mortality worldwide for 36 cancers in 185 countries. CA A Cancer J Clinicians 74, 229–263 (2024).

14. Cui, B. et al. Targeting ROR1 inhibits epithelial-mesenchymal transition and metastasis. Cancer Research 73, 3649–3660 (2013).

15. Li, C. et al. A ROR1-HER3-lncRNA signalling axis modulates the hippo-YAP pathway to regulate bone metastasis. Nat Cell Biol 19, 106–119 (2017).

16. Cao, J. et al. Twist promotes tumor metastasis in basal-like breast cancer by transcriptionally upregulating ROR1. Theranostics 8, 2739–2751 (2018).

17. Lu, D. et al. Structure-based discovery of quinazolin-2-amine derivatives as potent ROR1 pseudokinase inhibitors with in vitro and in vivo efficacy against triple-negative breast cancer. J. Med. Chem. 68, 16138–16171 (2025).

18. Shi, C. et al. Deep-learning-driven discovery of SN3–1, a potent NLRP3 inhibitor with therapeutic potential for inflammatory diseases. J. Med. Chem. 67, 17833–17854 (2024).

19. Wang, Y. et al. TF3P: Three-dimensional force fields fingerprint learned by deep capsular network. J. Chem. Inf. Model. 60, 2754–2765 (2020).

20. Jiang, K. et al. Rapid in silico directed evolution by a protein language model with EVOLVEpro. Science 387, eadr6006 (2025).

